# Distinct Neural Profiles of Frontoparietal Networks in Boys with ADHD and Boys with Persistent Depressive Disorder

**DOI:** 10.1101/2021.03.09.434662

**Authors:** Veronika Vilgis, Debbie Yee, Tim J. Silk, Alasdair Vance

**Affiliations:** Department of Paediatrics, University of Melbourne, Australia; Washington University in St. Louis, United States; Brown University, United States; Murdoch Children’s Research Institute, Australia; School of Psychology, Deakin University, Australia; Royal Children’s Hospital, Parkville, Australia

**Keywords:** fMRI, MVPA: working memory, ADHD, depression, children

## Abstract

Working memory deficits are common in attention-deficit/hyperactivity disorder (ADHD) and depression, two common neurodevelopmental disorders with overlapping cognitive profiles but distinct clinical presentation. Multivariate techniques have previously been utilized to understand working memory processes in functional brain networks in healthy adults, but have not yet been applied to investigate how working memory processes within the same networks differ within typical and atypical developing populations. We used multivariate pattern analysis (MVPA) to identify whether brain networks discriminated between spatial vs. verbal working memory processes in ADHD and Persistent Depressive Disorder (PDD). 36 male clinical participants and 19 typically developing (TD) boys participated in a fMRI scan while completing a verbal and a spatial working memory task. Within *a priori* functional brain networks (frontoparietal, default mode, salience) the TD group demonstrated differential response patterns to verbal and spatial working memory. The PDD group showed weaker differentiation than TD, with lower classification accuracies observed in primarily the left frontoparietal network. The neural profiles of the ADHD and PDD differed specifically in the SN where the ADHD group’s neural profile suggests significantly less specificity in neural representations of spatial and verbal working memory. We highlight within-group classification as an innovative tool for understanding the neural mechanisms of how cognitive processes may deviate in clinical disorders, an important intermediary step towards improving translational psychiatry.

## Introduction

Half of all mental illnesses begin prior to the age of 15 making them the leading cause of disability in young people worldwide (Kessler et al., 2007). Among the most common diagnosed neuropsychiatric conditions are attention-deficit/hyperactivity disorder (ADHD) and depressive disorders. ADHD and depression present distinctly in everyday life, yet research suggests some commonalities, as both disorders have been associated with impairments in attention (Kessler et al., 2007), executive function (Franklin et al., 2010; Martel, 2007; Martinussen et al., 2005; Matthews et al., 2008) and emotion regulation (Beauregard et al., 2006; Kovacs et al., 2008; Shaw et al., 2014). Additionally, neuroimaging studies have shown similar structural and functional anomalies in frontal, striatal, limbic, parietal and cerebellar brain regions in both disorders (Arnsten & Rubia, 2012; Cortese et al., 2012; Miller et al., 2015). Although evidence suggests that ADHD and depression potentially share some common cognitive and neural deficits, the extent to which such deficits are similar or distinct across both disorders remains unclear. Insight into how specific impairments (e.g., working memory) differ between ADHD and depression would improve characterization of key cognitive and neural processes disrupted and refine delineation in their underlying neurobiology.

One promising delineation between the working memory deficits often associated with ADHD and depression may be the specific type of working memory impaired. Children and adolescents with ADHD consistently demonstrate working memory deficits and tend to have greater impairments in processing spatial stimuli compared to verbal stimuli in these tasks (Martinussen et al., 2005; Sowerby et al., 2011; Willcutt et al., 2005). Conversely, although adults with depressive disorders also demonstrate working memory deficits (Gohier et al., 2009; Rose et al., 2006), they exhibit more persistent impairments in the verbal domain even after remission and independent of age (Gruber et al., 2011; Shilyansky et al., 2016). However, in contrast to the wealth of research implicating executive function deficits in children and adolescents with ADHD (Martel, 2007; Willcutt et al., 2005), much less is understood about the nature of executive function deficits in children and adolescents with depressive disorders (Vilgis et al., 2015). Nevertheless, the few extant studies have observed a deficit in working memory more often (Franklin et al., 2010; Günther et al., 2004; Klimkeit et al., 2005; Matthews et al., 2008; Vance & Winther, 2020) than not (Korhonen et al., 2002; Maalouf et al., 2011), though the extent to which verbal working memory deficits observed in adults also generalize to children and adolescents is unknown.

Both spatial and verbal working memory tasks consistently engage distinct neural representations in the frontoparietal network (FPN) (Daniel et al., 2016; Rottschy et al., 2012). In typically developing children and adults, the FPN exhibits hemispheric asymmetry, with greater right FPN activation for spatial working memory and left FPN activation for verbal working memory (Thomason et al., 2009). Children and adults with ADHD typically demonstrate less activation in the FPN during working memory tasks (Cortese et al., 2012), with hypoactivation over the right hemisphere compared to controls when processing visuospatial stimuli and bilateral prefrontal hypoactivation during a verbal task (Cubillo et al., 2014; Silk et al., 2005; Vance et al., 2007). Conversely, several studies in adults with depression have observed atypical activation of the left prefrontal cortex during working memory (Barch et al., 2003; Vasic et al., 2007; Walter et al., 2007), a pattern also observed in a pediatric sample of boys with Persistent Depressive Disorder (PDD) during a spatial task (Vilgis et al., 2014). Because disorder-specific hemispheric asymmetries exist independent of the type of stimulus material being processed, it is difficult to determine the extent to which atypical activation in ADHD and depression arises from dysregulated processing of a specific type of working memory (e.g., spatial vs. verbal) versus a more general neurological deficit associated with either disorder. Therefore, in order to develop greater understanding into how the neural patterns that underlie spatial and verbal working memory are altered in this network, it is necessary to perform a systematic comparison of how left and right FPN are disrupted in both working memory tasks in both ADHD and depression.

An important additional consideration is that networks that interact with the FPN may contribute to working memory deficits or play a compensatory role in the two disorders. In particular, the FPN interacts with the salience and default-mode network to support key working memory processes (Liang et al., 2016). The salience network (SN) plays a central role via continuously monitoring for endogenous and exogenous salient stimuli in order to facilitate the dynamic switching between internally directed, self-focused cognition carried out by the default-mode network (DMN) and externally directed cognition supported by the FPN (Menon & Uddin, 2010; Seeley, 2019). In children with ADHD, the absence of explicit task demands has been associated with weaker and more variable interconnectivity between FPN, SN, and DMN, including shorter and less persistent brain states (Cai et al., 2019). Similarly, in patients with depression, symptoms of rumination, emotional over-reactivity, and cognitive regulation of affective information have been associated with altered functional connectivity in the DMN, SN and FPN respectively (Hamilton et al., 2013; Manoliu et al., 2014). This compelling evidence suggests that the locus of working memory impairments in ADHD and depression likely arises from dysregulation of these core networks and their interactions. Yet, how neural patterns within each network support different types of working memory, and more critically, how these patterns putatively deviate in ADHD and depression, remains an enigma.

Thus, an outstanding question relates to understanding the precise mechanisms driving network-level differences in spatial and verbal working memory in ADHD and depression. Here, we present an innovative approach that combines network-based and multivariate analyses to investigate how neural patterns of spatial vs. verbal working memory putatively differ within clinical groups. Network-based approaches are uniquely powerful, as they provide a systematic approach for investigating neural systems underpinning cognitive function (Cole et al., 2014). Multivariate pattern analysis (MVPA) is a powerful machine learning method utilized to classify brain voxels to predict whether they carry information regarding distinct categories, tasks, or other cognitive states (Haxby et al., 2014; Mumford et al., 2012; Yang et al., 2012). Importantly, MVPA has been used to successfully dissociate between neural representations of cognitive tasks in prefrontal and parietal cortices (Etzel et al., 2016; Wisniewski et al., 2015; Woolgar et al., 2015). This precedent for utilizing MVPA to decode neural task representations strongly motivates the adaptation of this cutting-edge technique for examining networks that support working memory processes (e.g., FPN, SN, DMN).

Whereas MVPA has been previously adopted to investigate cognitive processes in typical development (Chow et al., 2018), such approaches have not yet been applied to understand how they diverge within clinical populations. In particular, MVPA may serve as a powerful tool that can systematically evaluate how distinctness of neural patterns underpinning different cognitive processes within the same brain networks may differ across typical vs atypical populations. Moreover, this data-driven method is aligned with computational psychiatry, an emerging field that promotes computationally rigorous approaches to improve the understanding, prediction, and treatment of mental disorders (Dwyer et al., 2018; Wiecki et al., 2015). Thus, combining MVPA with a network-based approach for clinical neuroimaging data provides a promising novel direction for improving mechanistic understanding of the distinctness of neural representations underlying spatial and verbal working memory tasks in ADHD and depression.

In the current study, we utilize MVPA to decode neural patterns underpinning spatial vs. verbal working memory in *a priori* brain networks (Hebart & Baker, 2018) within three different groups: typical development (TD), ADHD and PDD. We focus on PDD as a more chronic developmental disorder more similar to ADHD in its stability over time (versus major depressive disorder characterized by relatively shorter episodes of negative mood). An innovative feature of our within-group classification analysis entails neural decoding of spatial vs. verbal working memory processes in the same network. Given the known issues with how fMRI decoding results differ based upon the number of voxels in regions of interest (ROI) (Haynes, 2015), our approach of utilizing the same ROI for each decoding facilitates clear comparison of how neural representations of working memory processes are uniquely characterized within each clinical group. We predict distinct neural patterns in left and right FPN for spatial vs. verbal working memory processes in typical development. Moreover, we expect a double dissociation with an atypical pattern for the right but not left FPN in the ADHD group, and a deviant pattern for the left but not right FPN in the PDD group. Additionally, we examine neural patterns in the SN and DMN, two prominent networks known to interact with the FPN and previously identified as functioning atypically in ADHD and depression (Menon, 2011). Our objective was to leverage a novel technique to identify network response patterns within each developmental disorder, which may inform disorder-specific neural profiles underpinning well-known behavioral deficits in spatial and verbal working memory.

## Methods and Materials^1^

### Participants

39 male children and adolescents were recruited through an outpatient pediatric psychiatry clinic, met inclusion criteria for the study, and agreed to participate in the MRI scan. Of those, 17 fulfilled DSM-IV (American Psychiatric Association, 2000) diagnostic criteria for PDD (labelled as Dysthymic Disorder in DSM-IV) and 22 for a diagnosis of ADHD - Combined Type (i.e. at least six symptoms of inattentive and hyperactive/impulsive symptoms were met). All young people and their caregivers were interviewed using the Anxiety Disorders Interview Schedule for Children (Silverman et al., 2001). Caregivers also completed the Conners’ Parent Rating Scale - long version (Conners et al., 1998) and the Child Behavior Checklist (CBCL) (Achenbach, 1991) and the young person completed the Children’s Depression Inventory (Kovacs, 2003). An experienced child and adolescent psychiatrist (AV) confirmed each diagnosis. Participants were included if they were male, right-handed, free of metal implants (MRI compatibility) and with a full-scale IQ above 70. Exclusion criteria were the presence of an intellectual disability, learning disorder or known neurological or endocrine condition. Participants were also excluded if they had a previous diagnosis of an Autistic Spectrum Disorder, Bipolar Disorder or Psychotic Disorder. All clinical participants in the PDD and ADHD group had that disorder as primary diagnosis, but comorbidities of Oppositional Defiant Disorder, Conduct Disorder and anxiety disorders were permitted. Participants with ADHD (n=6) who were treated pharmacologically at time of testing withheld their stimulant medication at least 48h before participating in the MRI scan. Only one participant with PDD had been started on antidepressant medication prior to scanning. After excluding two ADHD participants due to excessive head motion and one PDD participant due to task performance, the final sample comprised 16 boys with PDD and 20 boys with ADHD. 19 typically developing participants (TD) and their caregivers were recruited through local schools. They completed the same questionnaires and semistructured clinical interviews as the clinical participants to ascertain normal psychological functioning. None of the TD met diagnostic criteria for any psychiatric diagnosis. An abbreviated IQ test was conducted with two subtests of the Verbal Comprehension Index (Similarities and Vocabulary) and two subtests of the Perceptual Reasoning Index (Block Design and Matrix Reasoning) (Wechsler, 2003) and scores were calculated as previously described (Crawford et al., 2010). All procedures were approved by the local Human Research Ethics Committee. All parents/guardians and young people gave informed written consent to participate in the study.

### fMRI Data Acquisition

Images were acquired on a 3T Siemens TIM Trio scanner (Siemens, Erlangen Germany) at the Royal Children’s Hospital, Melbourne. Participants lay supine with their head supported in a 32-channel head coil. High-resolution T1-weighted structural MRI images were acquired for each participant (TR=1900 ms, TE=2.24 s, FA= 90°, in-plane pixels=0.9 x 0.9 mm) prior to the functional scan. For functional imaging, T2*-weighted gradient-echo echo-planar images (EPI) were acquired (TR=2700 ms, TE=40 ms, FA=90°, 39 axial slices (co-planar with AC-PC) with 3.0 mm isotropic resolution). A total of 158 image volumes were acquired per 7 min 10 s sequence. This was run twice, one for each working memory task. Participants’ responses were recorded with a scanner compatible two button-box (Fibre-optic response pads, Current Designs, Philadelphia, PA).

### fMRI Task

A Sternberg type working memory task divided into verbal and spatial working memory components was adapted from Thomason et al. (2009). In the verbal task, participants viewed an array of capital letters, two in the low load condition and six in the high load condition. In the spatial task, either one or five black dots were presented randomly. After a 3000 ms delay, a single letter in lower case (verbal task) or a circle (spatial task) was presented for 1500 ms while participants indicated by button-press whether the probe matched the identity (verbal task) or location (spatial task) of the initial cues. Both tasks included control conditions that matched experimental conditions as much as possible (motor response, decision-making, visual stimuli, luminance, total trial length), except for the 3000 ms delay which was shortened to 100 ms. Probes matched the target 50% of the time. For each task, a total of 64 trials (32 experimental, 32 control) were presented in 16 pseudorandomly alternating blocks of four trials. Here, we combined high and low load conditions to optimize the beta estimates used in the MVPA analysis. Maximizing the number of blocks for within-subject classification improves accuracy via reducing overall noise of the beta estimates (Ku et al., 2008) and maximizes statistical power (Allefeld & Haynes, 2014). A more detailed task description is provided in the Supplementary Material and a schematic representation is shown in Figures 1a and 1b.

**Figure 1.**
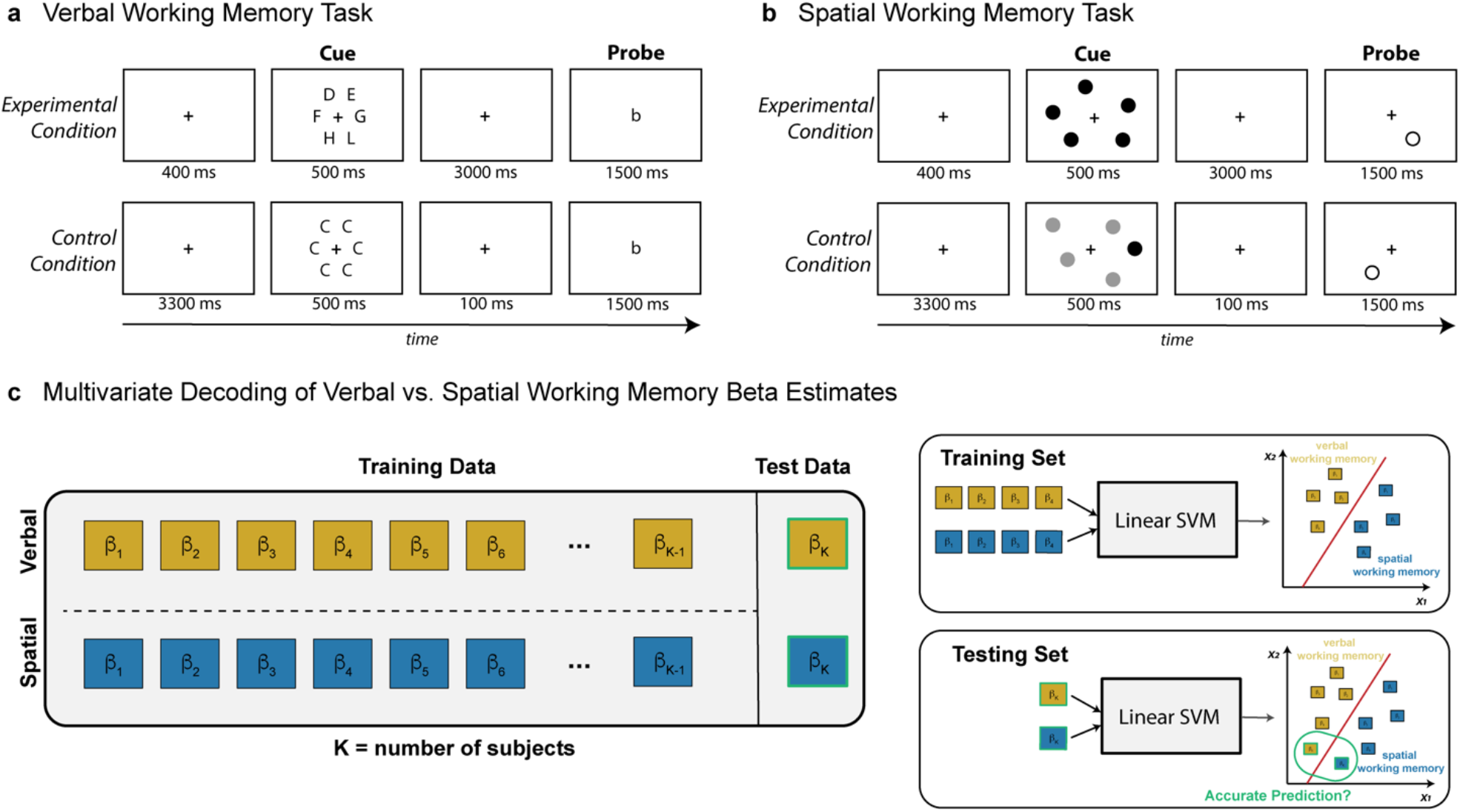
**a)** <u>Schematic of the Verbal Working Memory Task.</u> Participants were cued with an array of uppercase capital letters. After 3000 ms delay, a single lowercase letter was presented for 1500 ms while participants indicated by button-press whether the probe matched the identity of the initial cues. The control condition matched the experimental condition in as many elements as possible (motor response, decision-making, visual stimuli, luminance, total trial length), except for the 3000 ms delay which was shortened to 100 ms. **b)** <u>Schematic of the Spatial Working Memory</u> <u>Task.</u> This task was similar to the verbal task, except the cue was a set of black dots, and participants were tasked with indicating whether the circle probe matched the location of the initial task cues. **c)** <u>Multivariate Decoding of Spatial</u> <u>vs. Verbal Working Memory Beta Estimates.</u> A linear support vector machine (SVM) classifier with K-fold cross validation was implemented for each network within each group, with K equal to the number of subjects in each group. The classifier was trained on beta estimates for spatial and verbal working memory from K-1 subjects and tested on the remaining subject in that group. The process was repeated for the control conditions. Permutation tests (nsim=1000) evaluated whether observed classification accuracies for each network were statistically significant above chance.

### fMRI Preprocessing

Subject data was preprocessed using AFNI (https://afni.nimh.nih.gov/) (Cox, 1996). The EPI volumes were realigned, slice-timing corrected, registered to an MNI152 atlas, and normalized. EPI volumes were smoothed using a 4 mm FWHM Gaussian kernel to improve signal to noise ratio, which has been demonstrated in previous work (Gardumi et al., 2016; Woolgar et al., 2015). A general linear model was applied to each subject to extract block-level beta estimates for each of the four block types: spatial experimental, spatial control, verbal experimental and verbal control conditions. The design matrix was estimated using AFNI’s 3dDeconvolve function, and a generalized least squares time series fit with REML estimation of the temporal auto-correlation structure (3dREMLfit) was applied to estimate individual betas for the four regressors per subject. Each block lasted 21.6 seconds (8 TRs), and each run contained 16 blocks. These extracted beta estimates were fed into the linear Support Vector Machine (SVM) to classify spatial vs. verbal tasks for each of the 5 networks. Six motion parameters were included as GLM nuisance regressors. TRs (current and previous) were censored if the derivative values had a Euclidean norm above 3.5 mm.

To ensure the working memory tasks reliably activated frontoparietal regions in each clinical group, general linear tests were applied to the estimated beta weights to calculate task activation for spatial and verbal working memory independently for each participant. Within each group, AFNI’s 3dttest++ function was applied to generate statistical maps for the average of the spatial and working memory conditions. Significant clusters for each task condition and group (thresholded at p<.001, cluster corrected with threshold *α* = .01) are shown in Figure 5.

### ROI Masks

The ROI masks were selected from https://www.fmrib.ox.ac.uk/datasets/brainmap+rsns/ and have been described previously (Smith et al., 2009). We focused on the left and right FPN, key networks supporting working memory related activity and in accordance with our predictions that classification accuracy would differ between the groups given the previous literature. In addition, we included the DMN and the SN, as these networks dynamically interact with the FPN to support working memory. We also included the sensorimotor (SM) network as control network, where we do not expect significant differences between tasks or groups. All brainmaps were resampled to match the dimensions of the data (3×3×3 mm isotropic), thresholded at 3.5, and binarized to create masks for each of the networks. The ROIs are shown in Figure 3a.

**Figure 2:**
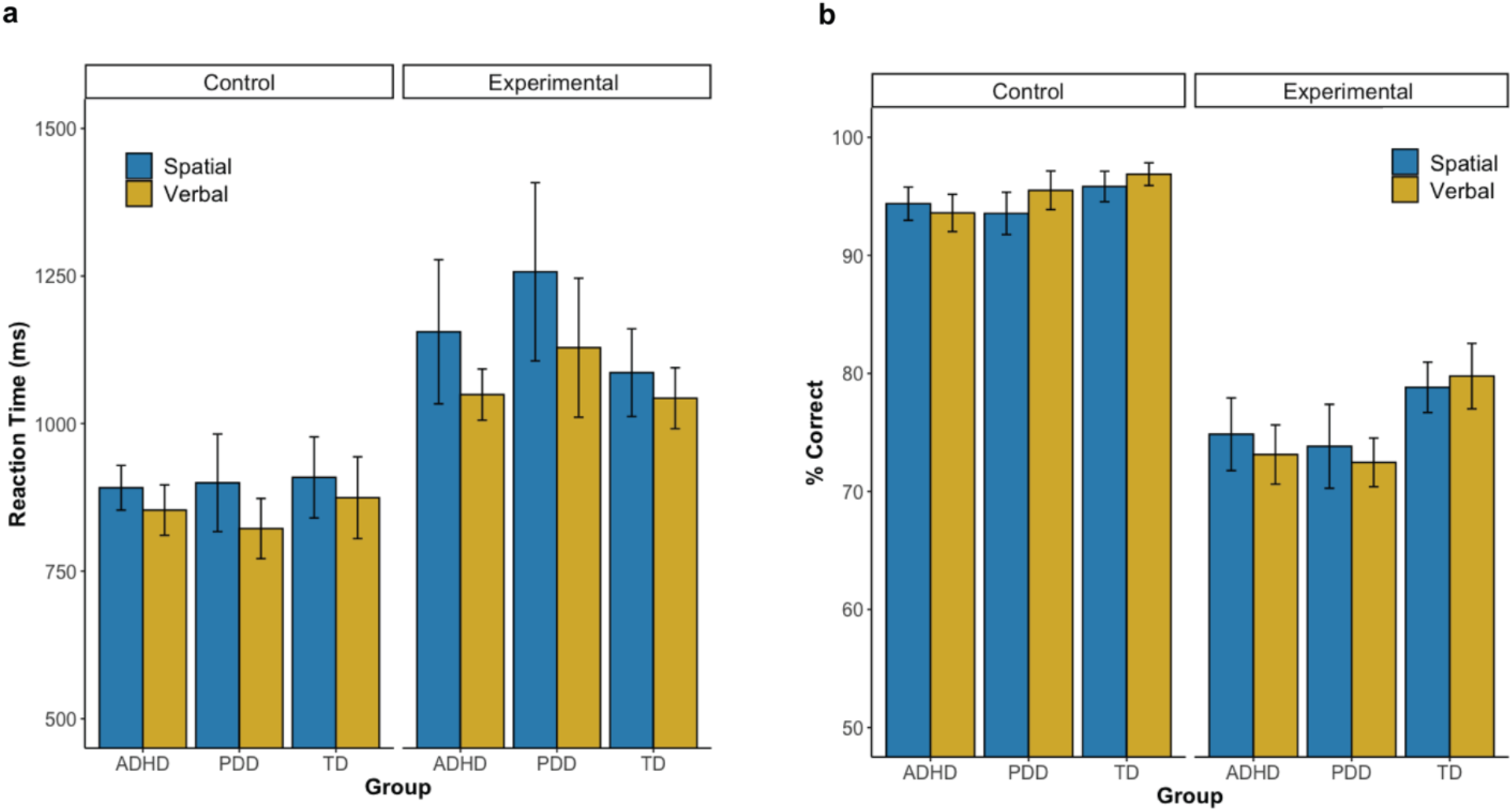
a) <u>Reaction Time by Experimental and Control Condition</u>. Participants were overall slower when performing the experimental conditions relative to the control conditions. Although RT performance was similar between verbal and spatial memory tasks, there was a slight trend towards faster RT in the verbal condition in the PDD group. For both plots, error bars indicate standard error. **b)** <u>Accuracy by Experimental and Control Condition.</u> Participants were overall less accurate when performing the experimental conditions relative to the control conditions. For each group, the accuracy did not differ between verbal and spatial memory task components.

**Figure 3.**
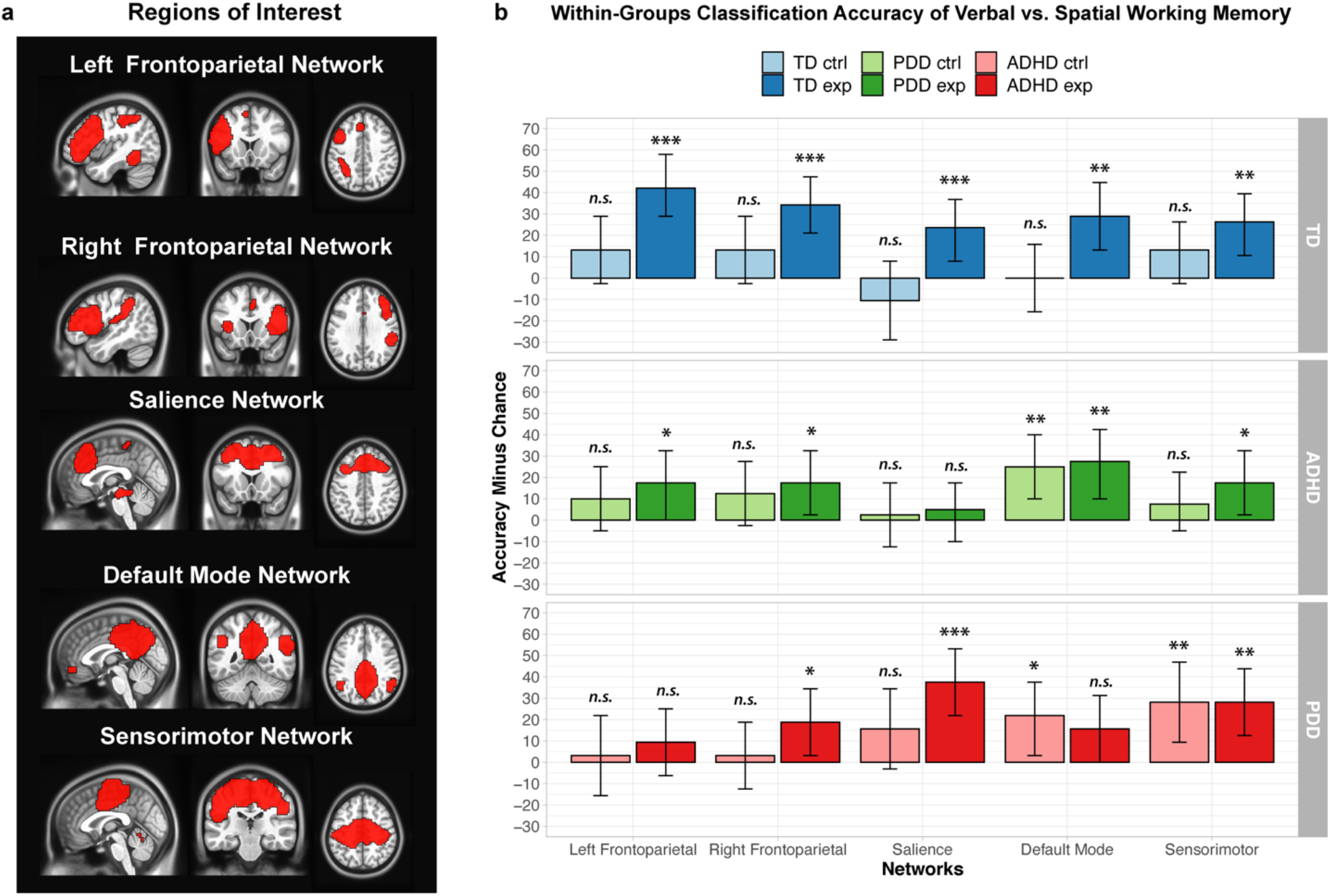
a) <u>Regions of Interest</u>. Left and Right Frontoparietal Network (LFPN and RFPN) forming as the primary ROIs for working memory function, followed by the Salience Network (SN) and Default Mode Network (DMN) known to interact with the FPN. The sensorimotor network (SM) served as control ROI. b) <u>Within-Groups</u> <u>Classification Accuracy of Spatial vs. Verbal Working Memory for the five selected networks.</u> Here, the classification accuracy indicates the distinctness of spatial vs. verbal working memory task representations within each ROI. Importantly, utilization of the same ROI for each decoding facilitates clear comparison of how neural representations of working memory tasks may differ across clinical groups. High classification accuracies are observed across all networks but with group specific patterns. The TD group shows distinguishable patterns of verbal and spatial working memory processes in all networks. Classification accuracies are significantly high for the right FPN, salience and sensorimotor networks but not the left FPN or DMN in the PDD group. For the ADHD group we observed significant classification accuracies only in the DMN with a trend for the left FPN. Error bars represent 90% confidence intervals, and statistical significance is listed for one-tailed permutation tests: *p<.05; **p<.01; ***p<.001. Permutation distributions for each ROI, condition, and group are illustrated in Figure S2 in the Supplementary Material.

### Multivariate Pattern Analysis (MVPA)

Multivariate pattern analysis (MVPA) was performed on each of the 5 networks (left and right FPN, SN, DMN, SM) using the Decoding Toolbox (Hebart et al., 2015), and custom MATLAB code. All classifications implemented a linear SVM classifier using a classification kernel from libsvm package with a fixed cost parameter (c=1). K-fold cross-validation was performed for each group using a leave-one-run-out protocol, with K equal to the number of subjects in each group. Here, the classifier was trained on data from K-1 subjects (per group) and tested on the left out remaining subject in that group. The beta-estimates were used to train and test our classifier (Pereira et al., 2009) on spatial vs. verbal working memory (See Figure 1c). Critically, the utilization of K-fold cross-validation allows for better generalization of the classification model to novel datasets (e.g., less overfitting to the dataset).

The linear SVM classifier generates signal detection metrics that could be used to compare the classification accuracy (e.g., the proportion of correctly classified observations), as well as additional performance metrics such as the model’s sensitivity (e.g., true positive rate) and model’s specificity (e.g., true negative rate). In the current study, we summarize these signal detection metrics into area under the curve (AUC) parameter, which allows us to incorporate both sensitivity and specificity measures to evaluate the intrinsic accuracy of the diagnostic test and compare classifier performance measures between networks and groups (W. Zhu et al., 2010). Greater detail of these performance metrics is provided in the Supplemental Material. Permutation tests (nsim=1000) evaluated whether the observed cross-validated classification accuracies for each network were statistically significant above chance (Etzel & Braver, 2013; Hebart et al., 2015). An important feature of such non-parametric permutation tests is that they allow for estimation of the statistical significance by estimating the probability of obtaining an observed classifier performance under the null hypothesis, based on the estimated empirical cumulative distribution of classifier error (Golland and Fischl, 2003). Critically, these tests provide stable empirical p-values based on these distributions (Ojala & Garriga, 2010). The procedure was repeated for the spatial versus verbal control conditions.

## Results

### Clinical and Behavioral Results

We first characterized how clinical symptoms differed at the group level, which would inform subsequent within-group differences. Both clinical groups exhibited greater levels of externalizing and internalizing symptoms compared to the TD group and lower scores on IQ measures. As expected, both clinical groups differed from each other in scores of internalizing symptoms (PDD > ADHD) and related global ADHD and restless symptoms (ADHD > PDD). Summary statistics of the sample characteristics are presented in Table 1.

**Table 1:**
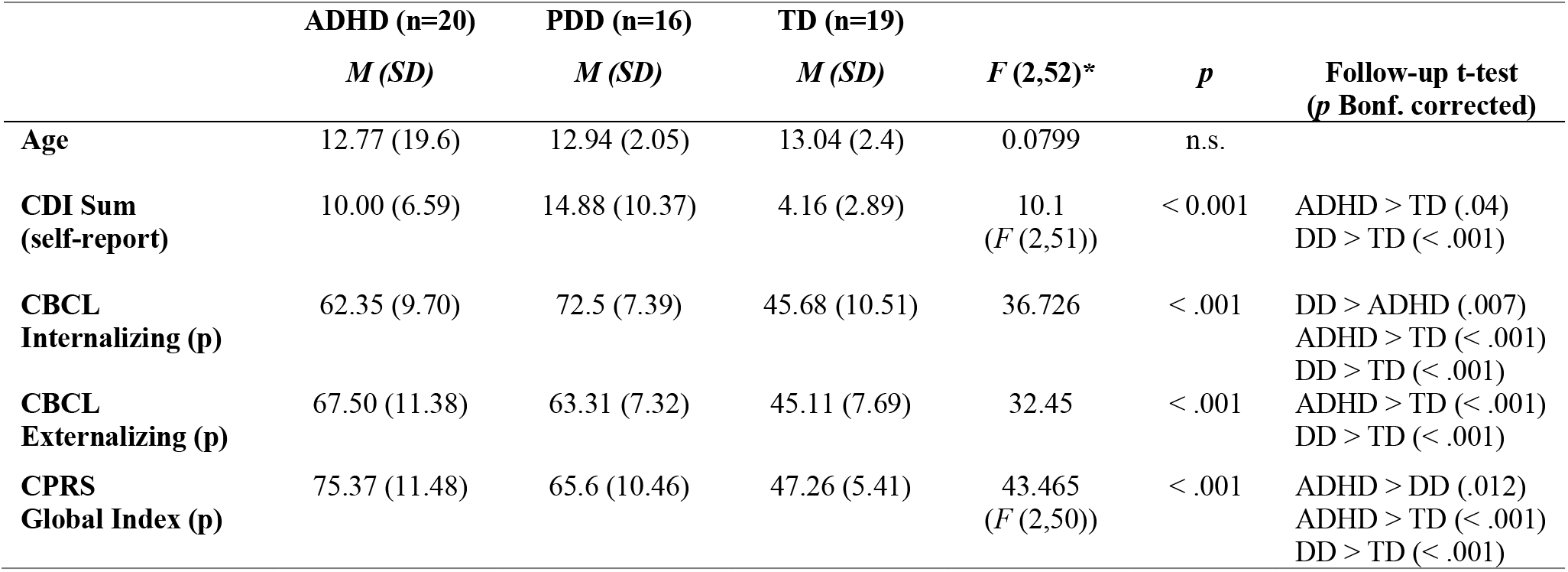
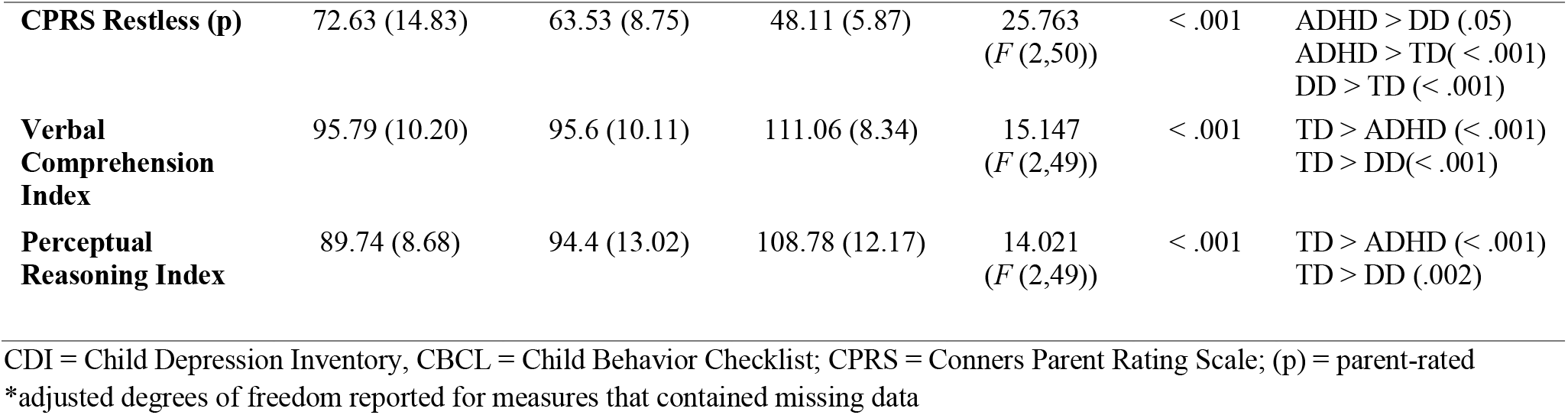
Means (M) and standard deviations (SD) of demographic and clinical variables for each group.

With regard to the behavioral performance on the two working memory tasks, within each group, reaction times (RT) and accuracy did not differ between the verbal and spatial task components (Accuracy: PDD *t*(15)=0.37, *p*=0.717; ADHD *t*(19)=0.61, *p*=0.547; TD *t*(17)=-0.31, *p*=0.758; RT: PDD *t*(15)=-1.21, *p*=0.246; ADHD *t*(19)=-0.97, *p*=0.342; TD *t*(18)=-1. 27, *p*=0.222). In the control conditions, behavioral performance was similar between both tasks, though there was a trend towards faster reaction times in the verbal than spatial control condition in the PDD group (Accuracy: PDD *t*(15)=0.92, *p*=0.370; ADHD *t*(19)=-0.50, *p*=0.624; TD *t*(17)=0.89, *p*= 0.385; RT: PDD *t*(15)=-2.20, *p*=0.044*, ADHD *t*(19)= -1.43; *p*=0.169; TD *t*(18)=- 1.62, *p*=0.123). Task performance measures are illustrated in Figure 2 and provided in Table S1 in the Supplementary Material. Additionally, motion did not differ between verbal and spatial working memory conditions nor control conditions (See Table S2 in Supplementary Material).

### Multivariate Decoding Results (MVPA)

We applied a linear SVM to *a priori* brain networks and identified disorder-specific classification accuracies that indicate the distinctness of spatial vs. verbal working memory task representations within each network. These networks are shown in Figure 3a. Table 2 presents the cross-validated classification accuracy minus chance (chance = 50%) from the decoding analyses for each ROI. Our primary focus was on the left and right FPN within each group. For both networks, we expected to observe significant classification accuracies in the TD group indicative of differential neural representations of verbal and spatial working memory. Indeed, the TD group showed significantly above chance classification accuracies within both the left (*p*=.001) and right (*p*<.001) FPN. For the ADHD group, significant classification accuracies were observed within the left FPN (*p*= .034) and right FPN (*p*=.035). For the PDD group, classification accuracy was not significant in the left FPN (*p*=.223) but was statistically significant within the right FPN (*p*=.030).

**Table 2:**
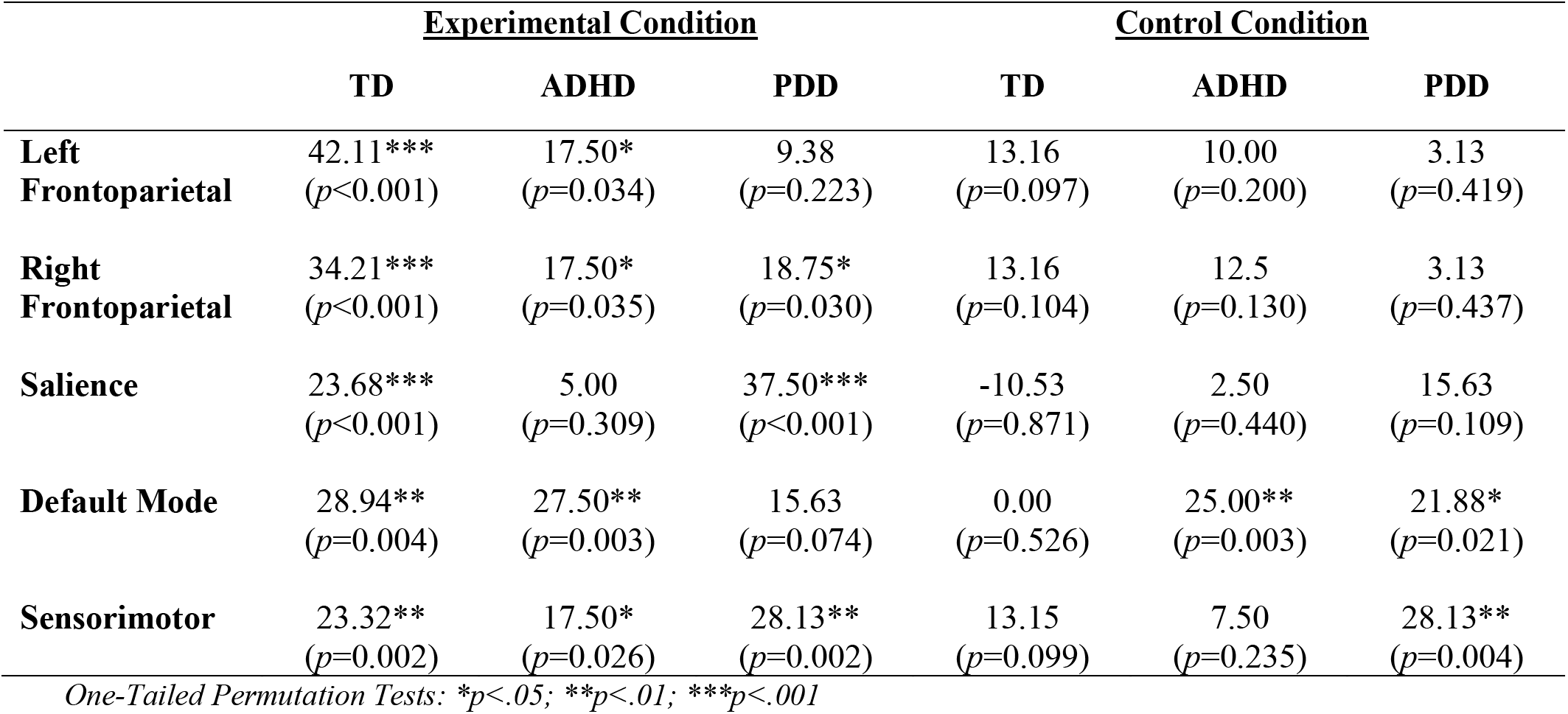
Within Groups Classification Accuracy Minus Chance for each group and region of interest separately for the experimental (working memory) and control condition.

Additionally, we included the SN and DMN as networks of interest due to their known interactions with the FPN and prior work indicating aberrant within and across network functions in both ADHD and PDD. For the TD group, both the SN (*p*<.001) and DMN (*p*=.004) exhibited significant classification accuracies. For the PDD group, classification accuracy was significant for the SN (*p*<.001) but not the DMN (*p*=.074). Conversely, for the ADHD group, classification accuracy was significant for the DMN (*p*=.003) but not for the SN (*p*=.309).

We also examined the classification accuracies of the spatial vs. verbal control conditions (trials that matched the experimental working memory trials in all aspects with the exception of the delay period, thus removing the memory component of the task). None of the classification accuracies for these control conditions were significant in the TD group, suggesting that spatial and verbal information does not *per se* elicit differential processing in the right and left FPN, DMN and SN. In the PDD and ADHD groups, we observed the same pattern as in the TD group in the left and right FPN and SN. However, the DMN showed significant differential processing of verbal and spatial material in the control condition for both the PDD group (*p*=.021) and ADHD group (*p*=.003).

Lastly, we examined the SM as a control network, and predicted that classification accuracies would be similar in all three groups with little differentiation between verbal and spatial processes. Unexpectedly, in all three groups classification accuracy was significantly high for the working memory trials. For the control trials, we did not observe significant differentiation for TD and ADHD groups, but observed significant classification accuracy for the PDD group (*p*=.004). Figure 3b displays the classification accuracy minus chance across ROIs, delineated for each group.

### ROC and AUC Metrics for Comparison Between Groups

In order to facilitate quantitative comparison of the classification model performance between ROI networks and groups, we utilized signal detection metrics and estimated the receiver operator characteristic (ROC) curve and area under the ROC curve (AUC) metrics. The **ROC** curve reflects a combination of the model’s ***sensitivity*** (i.e., proportion of true positive assessments) and ***specificity*** (i.e., proportion of true negative assessments), which can be used to compute an **AUC** measure of classifier performance for each network and each group. Critically, as the AUC measure considers the tradeoff between true positive rate and false positive rate, it provides a more comprehensive measure to compare classifier performance in each network between groups. Greater detail of how these signal detection metrics are derived and calculated is described in the Supplementary Material, and a table of all of the metrics is provided (Table S3).

Comparing across groups, the AUC was numerically higher in the TD group compared to PDD and ADHD groups (AUC_TD_ = 97.51, AUC_PDD_ = 66.02, AUC_ADHD_ = 73.50) for the left FPN, and permutation tests revealed significant group differences between TD vs PDD groups (AUC_TD-PDD_ = 31.49, *p*=.027) but not between ADHD and PDD groups (AUC_ADHD-PDD_ = 7.48, *p*=.330). Although the AUC difference between TD and ADHD groups was substantial, this difference did not reach statistical significance (AUC_TD-ADHD_ = 24.00, *p*=.073). In the right FPN, the AUC in the TD group was also numerically higher compared to both groups (AUC_TD_ = 85.32, AUC_PDD_ = 73.83, AUC_ADHD_ = 65.25), though these differences did not reach statistical significance between TD and PDD groups (AUC_TD-PDD_ = 11.49, *p*=.191), TD and ADHD groups (AUC_TD-ADHD_ = 20.07, *p*=.093), or ADHD and PDD groups (AUC_ADHD-PDD_ =-8.58, *p*=.326).

Interestingly, the pattern of ROC curves was different for both the SN and the DMN. Specifically, within the SN, the AUC was highest for the PDD group (AUC_PDD_ = 93.75), followed by the TD group (AUC_TD_ = 77.84), and close to chance for the ADHD group (AUC_ADHD_ = 56.50). The AUC difference was statistically significant between ADHD and PDD groups (AUC_ADHD-DD_ = -37.25, *p*=.009), but did not reach threshold for significance between TD and ADHD groups (AUC_PDD-TD_ = 21.34, *p*=.073) nor TD vs PDD groups (AUC_PDD-TD_ = -15.91, *p*=.165). Within the DMN, the AUC was similar for both TD and ADHD groups (AUC_TD_ = 84.49, AUC_ADHD_ = 84.00), both of which were numerically higher than the AUC for the PDD group (AUC_PDD_ = 74.22). None of these group differences were however, significantly different (AUC_TD-ADHD_ = .49, *p*=.535, AUC_TD-PDD_ = 10.27, *p*=.280, AUC_ADHD-PDD_ = 9.78, *p*=.242). Finally, within the SM network, the AUC is similar for both TD and PDD groups (AUC_TD_ = 80.61, AUC_PDD_ = 83.59), both of which were higher than the ADHD group (AUC_ADHD_ = 68.25). There were no significant group differences observed for the SM network (AUC_TD-ADHD_ = 12.36, *p*=.226, AUC_TD-PDD_ = -2.98, *p*=.431, AUC_ADHD-PDD_ = -15.34, *p*=.164). Figure 4a plots each group’s ROC curve by network and 4b provides corresponding group AUC metrics. Permutation distributions of the group differences are detailed in Table S4 and Figure S3 in the supplemental material.

**Figure 4.**
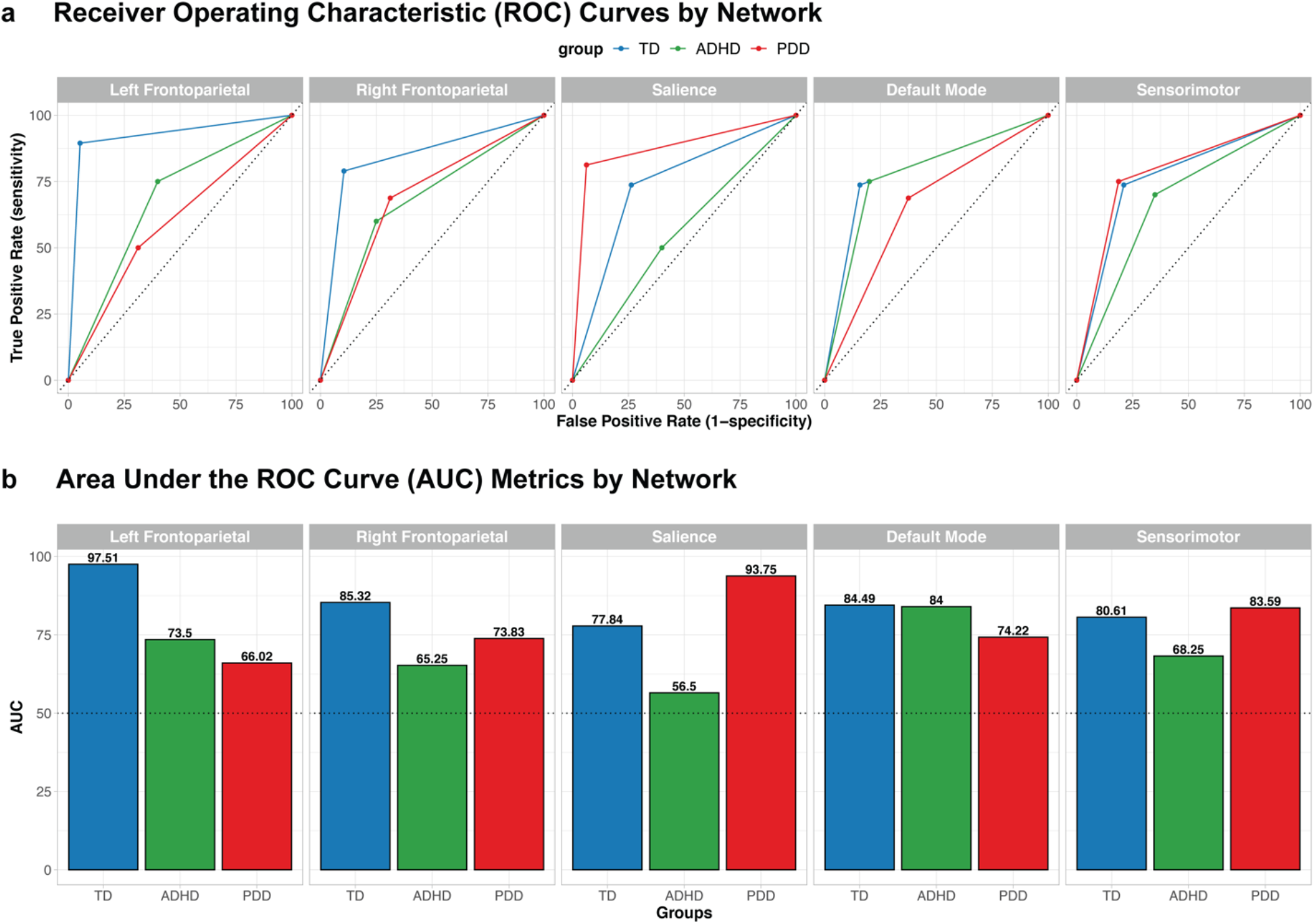
a) <u>Receiver Operating Characteristic (ROC) Curves by ROI Network and Group.</u> These empirical ROC curves reflect the combination of the linear SVM’s sensitivity (true positive rate) and specificity (false positive rate), which we combine to compute an area under the ROC curve (AUC) measure of classifier performance for the control condition. The colored lines indicate the ROC curves for TD (blue), ADHD (green), and PDD (red) groups. The black diagonal dotted line indicates chance performance. b) <u>Area Under</u> <u>the ROC Curve (AUC) By ROI Network and Group</u>. Notably, the AUC measures are highest for the TD group in both left and right Frontoparietal Networks (FPN), whereas the AUC is highest for the PDD group in the Salience Network (SN). The highest AUC measures in the Default Mode Network (DMN) are similar for both TD and ADHD groups. The black dotted line indicates chance performance.

### Whole Brain Univariate Results for Spatial and Verbal Working Memory Tasks

In order to verify that the fMRI task reliably activated key working memory related regions, we averaged whole-brain activations for spatial and verbal working memory experimental conditions for each group (TD, ADHD, PDD). Our results revealed that all groups show expected activation in left and right FPN and SN, as well as expected deactivation in the DMN. Importantly, similar activation of these networks across all groups provide evidence that these key working memory networks are involved in both spatial and verbal tasks. Averaged beta estimations for each group are shown in Figure 5, and MNI coordinates of peak activations for all conditions are shown in Table S5 in the Supplemental Material.

**Figure 5:**
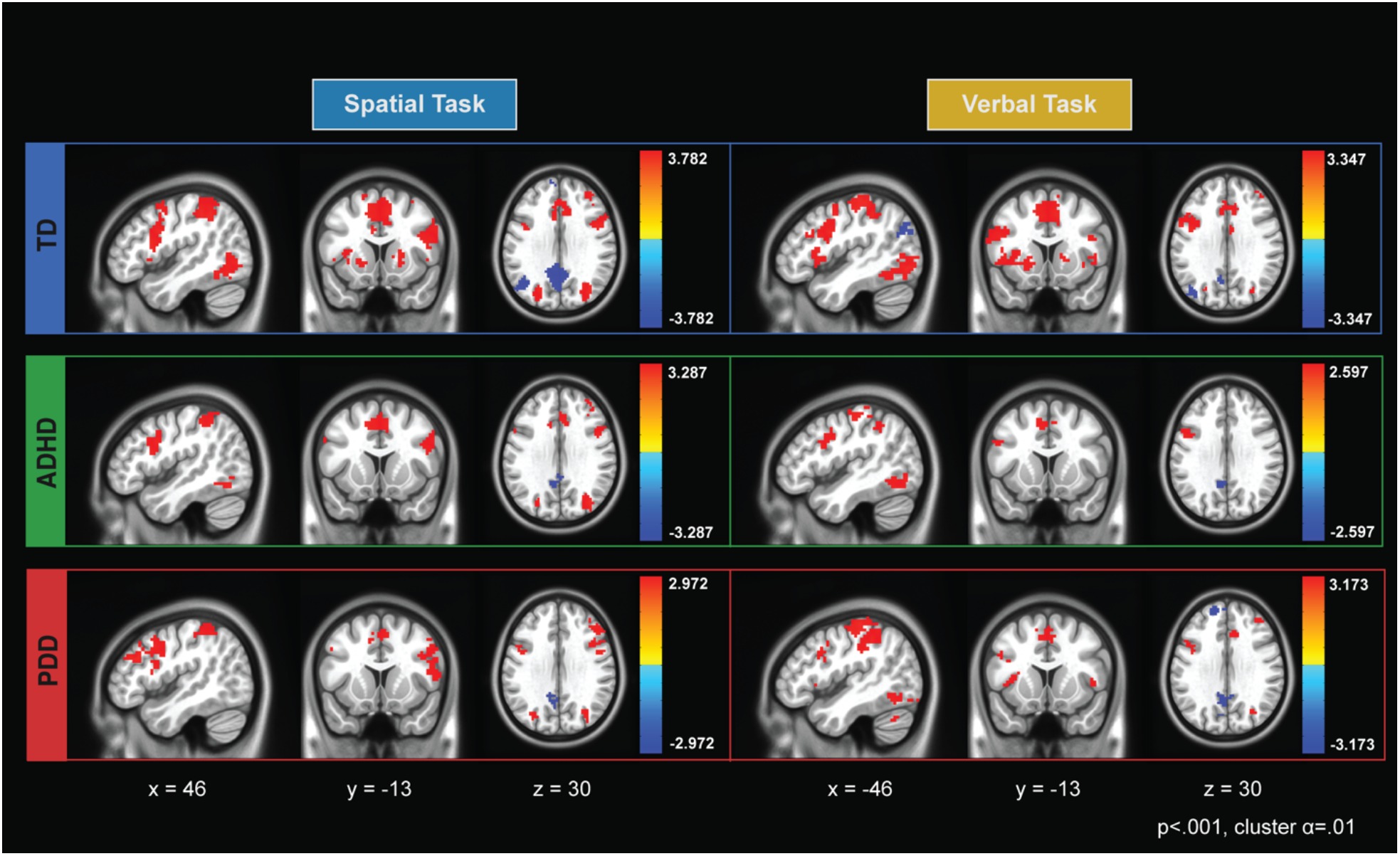
Whole-brain activations for the spatial and verbal working memory experimental condition in typical development, ADHD, and PDD groups. These results revealed all groups show expected activation in key working memory related regions (e.g., left and right frontoparietal networks, salience network) and expected deactivation in default mode network for both spatial and verbal tasks. Importantly, these data show consistent activation in these networks across all groups for both spatial and verbal tasks, which markedly deviate from the neural profiles of clinical groups that were established from the MVPA decoding analyses. Significant clusters were thresholded at p<.001 and cluster corrected with a threshold of *α* = .01. The colors correspond to t-values, with yellow-red associated with positive values and blues associated with negative values.

## Discussion

In this study, we identified distinct neural profiles for spatial versus verbal working memory processes in frontoparietal, salience, and default mode networks in boys with ADHD, boys with PDD, and typically developing male youth. We used a novel approach that combined MVPA and network-based analyses to compare multivariate neural representations of verbal and spatial working memory within each group. Our approach revealed clear distinguishable neural patterns in the TD group in *a priori* networks not observed in either clinical group. The PDD group differentially processed verbal and spatial material in right but not left FPN while classification accuracies in the ADHD group showed significant differential processing in both networks. Partly in line with our predictions, we observe a higher AUC value in the left FPN and a lower value for the right FPN in the ADHD compared to the PDD group. However, significant group differences in classifier performance were detected only between the TD and PDD groups for the left FPN. In addition, a significant difference in classifier performance for the SN also revealed differing neural profiles of the clinical groups with distinct processing of spatial vs. verbal working memory in PDD but not in ADHD.

The TD group’s neural profile shows clear distinctions between verbal and spatial working memory processes across all networks. In contrast, no differential processing of verbal and spatial material was observed in control conditions. The highest classification accuracies were observed in the left and right FPN, demonstrating known lateralized processes such as verbal rehearsal (left > right) (Nagel et al., 2013) or spatial information processing (right > left) (Ray et al., 2008). The significant classification accuracies in the DMN may be attributed to greater involvement of posterior DMN regions in mental imagery (i.e., precuneus), supporting retention of non-verbal material (Cavanna & Trimble, 2006). Lastly, the significant distinctiveness found in the SN likely suggests that one type of information, verbal or spatial, may be more or less salient in the context of working memory.

The PDD group revealed a neural profile of significant differentiation between verbal and spatial working memory in the right FPN and SN, and non-significant differentiation in the left FPN and DMN. Interestingly, both left and right FPN show no differential processing of verbal and spatial material in the control condition, but only the right FPN successfully differentiates during the working memory condition. This pattern suggests that pediatric PDD may be associated with a failure to trigger specialized processing patterns in the left FPN during working memory, consistent with prior work implicating aberrant functioning of left FPN during working memory tasks in adults with depression (Vasic et al., 2007; Walter et al., 2007). Differentiation between verbal and spatial processes in the DMN was detected during the control condition and not during the experimental condition, a pattern opposite to the one observed in the TD group. Studies showing opposing activation and functional connectivity patterns of the FPN, DMN and SN between adult patients with depression and non-depressed individuals have reported compensatory mechanism in these networks which support successful working memory performance in patients but not in controls and vice versa (Albert et al., 2019; Harvey et al., 2005). The unique neural profile seen in the PDD group therefore likely reflects both pathophysiological mechanisms in some networks such as the left FPN as well as compensatory mechanisms in DMN or SN.

With the exception of the SN, the ADHD group’s neural profile indicates differentiation of verbal and spatial working memory processes in all networks. The results for the SN suggest saliency detection and dynamic switching between networks are not sufficiently different between the verbal and spatial tasks. In contrast, significant classification in the DMN in both the experimental and the control condition suggests processing differences between the two tasks that are not specific to working memory. DMN altered within and between network connectivity as well as increased activity in this network during cognitive tasks is a common finding in patients with ADHD, and has been associated with excessive mind-wandering, attentional lapses and impaired intertemporal decision making (Bozhilova et al., 2018; Castellanos et al., 2009; Sonuga-Barke et al., 2016; Sonuga-Barke & Castellanos, 2007). Our results suggest that the neural response patterns in the DMN are sensitive to the task context and behave differently whether verbal or spatial stimuli are presented. As the FPN, SN and DMN are tightly connected and work together for optimal cognitive performance (Menon, 2011), suboptimal function in just one network likely has effects on the functioning of the other. Future studies may therefore want to examine more closely the between network dynamics that result in deviant neural processing in the SN and DMN observed here to inform neurobiological mechanisms underpinning working memory deficits in pediatric ADHD.

Additionally, we included the sensorimotor network as a control region of interest and expected comparable classification accuracies across groups. For the working memory conditions, all three groups showed significant differential processing of verbal and spatial tasks, with similar AUCs for the TD and PDD group and with a lower but not significantly different AUC for the ADHD group. Unexpectedly, for the control condition the PDD group also showed high classification accuracy which was not observed in the two other groups. Although the tasks were closely matched, and we did not observe significant behavioral differences in reaction times and accuracy across tasks, we cannot exclude the possibility that systematic differences may have contributed to the distinct neural patterns observed in the SM and other networks. Individual differences in dynamic functional connectivity of this network (and the FPN) have previously been shown to correlate negatively with working memory capacity and accuracy (Zhu et al., 2021). Furthermore, parts of the sensorimotor network support verbal and spatial maintenance processes (Smith & Jonides, 1999) and task-based fMRI studies have linked activity in this network to the number of stimuli held in mind (Kirschen et al., 2005; Marvel & Desmond, 2010) a factor that differed between the verbal and spatial tasks used here. Therefore, given these known contributions of the SM network to working memory processes, it may not have been an adequate network to use as control region.

Lastly, in addition to our within-group classification analyses, we also examined signal detection metrics from the linear SVM to compare classifier performance across groups for the experimental condition. In particular, comparison of AUC metrics revealed that the classifier performed better in the TD group than in the PDD group for the left FPN. Overall, our results are consistent with the large body of research that has shown differences in the neural underpinnings between patient groups diagnosed with clinical depression and non-clinical groups during working memory tasks (e.g., Matsuo et al., 2007; Vilgis et al., 2014; Walter et al., 2007; Wolf et al., 2009).While we found no significant group differences between the ADHD and TD group for three networks (right and left FPN and SN), the TD group had consistently larger classification accuracies. The absence of any significant group differences may be due to our study being underpowered. Additionally, classifier accuracy differed significantly between the two clinical groups for the SN. These results provide novel information regarding disorder-specific neural profiles during working memory, which have important implications given the high comorbidity of the two disorders. Although both disorders have repeatedly been associated with abnormal function and connectivity in the core networks investigated in the current study, we demonstrate not only differences between the clinical groups and typically developing youth but also between the clinical samples. Future studies should aim to delineate how divergent network function during working memory contributes to different disorders or even distinct psychiatric symptom dimensions, which may be informative for elucidating the potential behavioral or cognitive factors that drive unique or divergent neural profiles of youth with ADHD and youth with depressive disorders.

We acknowledge important limitations, including the lack of spatial specificity (i.e., which voxels within each network are most distinct) or whether deviant responses to either the verbal or spatial task (or both) contributed to significant differentiation of neural patterns. While MVPA can discriminate between neural patterns underlying different cognitive processes within a specific population, more work needs to be done to explicitly compare groups. A logical next step would be to test the degree to which models trained on one group can accurately classify task performance in a different clinical group, or apply complementary multivariate approaches that could determine shared and unique neural response patterns across clinical diagnoses (e.g., representational similarity analysis; Kriegeskorte & Diedrichsen, 2019). Additionally, although we opted to collapse across the load-component of the two tasks to increase the reliability of the beta-estimates for the classification analysis, more systematic evaluation of working memory load in subsequent studies may help identify additional clinically relevant neural processing patterns in typical and atypical development. Nevertheless, by separately examining the working memory and control conditions we were able to show clear distinctions between more cognitively demanding and simple perceptual processes, respectively. Another technical consideration relates the limited temporal resolution of our beta estimates, due to the relatively slow TR (2.7s) and blocked design of the working memory paradigm. Future studies may opt to capitalize upon recent advances in multiband acquisition sequences (Bhandari et al., 2020; Risk et al., 2021) that will allow for shorter TR measurements and improve temporal specificity of beta estimates used for MVPA or other types of multivariate analyses. Notably, although our sample well-matched across groups (thus making it well-suited for multivariate pattern classification), it was relatively small and focused on only male participants within a limited age range, which may restrict the generalizability of the results. It is also not clear to what extent our results may apply to adults with ADHD or PDD. However, up to 94% of individuals with early onset PDD (i.e., before the age of 21) report a subsequent a major depressive episode at some point in their life (Klein et al., 2000) suggesting neural alterations early on may constitute a vulnerability for later depression and likely stay constant or worsen with age. For those with ADHD, between 50% to 60% show persistence of symptoms into adulthood (Roy et al., 2016), particularly symptoms of inattention that are more closely linked to executive function deficits such as working memory. As such, while studies in adults are necessary to confirm the transferability of our findings, we believe the findings and methodology equally applies to adults with the disorder. Finally, although our small sample is well characterized clinically, it is not well-suited for diagnostic (i.e., between-subjects) classification. Future studies could use classification approaches to predict whether dissociable network patterns predict clinical diagnoses (Nielsen et al., 2020; Saeed, 2018; Sundermann et al., 2014) or subtypes within diagnosis (Fair et al., 2013). Such investigations could focus on transdiagnostic measures, which entails predicting symptom dimensions rather than categorical diagnoses, as proposed by the Research Domain Criteria Framework (Buckholtz & Meyer-Lindenberg, 2012; Cuthbert & Insel, 2013; Parkes et al., 2020).

## Conclusion

In conclusion, we highlight within-group classification as an innovative approach to identify the distinctness of neural response patterns in *a priori* brain networks across two neurodevelopmental disorders. We found group-specific neural profiles comprising network-specific spatial vs. verbal working memory processes. The neural profile of the TD group is characterized by distinct neural response patterns when retaining spatial vs. verbal information in working memory, suggesting greater specialization for and sensitivity to task demands in several key networks supporting essential cognitive, emotional and social functions. The neural profiles of both clinical groups deviate from the control group, hinting at neural representations putatively less responsive to varying task demands. Identifying group-specific neural profiles of essential cognitive processes in health and disease is a much-needed intermediary step to advance translational psychiatry and to help inform dysregulated brain mechanisms (Huys et al., 2016; Woo et al., 2017). Critically, we demonstrate that data-driven multivariate approaches can help gain a more nuanced understanding of how brain networks underpinning cognitive processes may differ within clinical diagnoses.

## Acknowledgments

This research was conducted within the Developmental Imaging research group, Murdoch Children’s Research Institute and the Academic Child Psychiatry Unit, Department of Paediatrics, The University of Melbourne. Royal Children’s Hospital, Melbourne, Victoria. It was supported by the Murdoch Children’s Research Institute, the Royal Children’s Hospital, The Royal Children’s Hospital Foundation, Department of Paediatrics, The University of Melbourne, and the Victorian Government’s Operational Infrastructure Support Program. TS was supported by a NHMRC Career Development Award. DY was supported by the National Institutes of Health NRSA Fellowship F31-DA042574. We would like to thank all children and families who participated in this research. We would also like to thank Dr. Lauren Richmond and Dr. Alec Solway for helpful comments and suggestions on the manuscript. DY and VV would also like to thank the Kavli Foundation for enabling participation in their Summer School in Cognitive Neuroscience (2016, Santa Barbara, USA), where this collaboration originated.

## Open Practices Statement

The data and materials for all experiments are available on our OSF repository: https://osf.io/a5349/. The experiment was not preregistered.

## Footnotes

1 All task scripts and analysis scripts are uploaded on our OSF repository: https://osf.io/a5349/

