## Supplementary Material for "Distinct Neural Profiles of Frontoparietal Networks in Boys with ADHD and Boys with Persistent Depressive Disorder"

**Method**

*fMRI task*

A Sternberg type working memory task divided into verbal and spatial working memory components was adapted from Thomason et al. (2009). Each trial began with a central fixation cross displayed for 400 ms. In the verbal task participants were presented with an array of capital letters, two in the low load condition and six in the high load condition for a duration of 500 ms. After a 3000 ms delay a single letter in lower case was presented for 1500 ms and participants indicated by button-press whether the letter was present in the previously shown array. In the verbal control condition only one type of letter was shown, two in the low and six in the high load condition and the 3000 ms delay was shortened to a 100 ms delay. Only consonants were used and letters were not repeated if they appeared in the previous two trials. In the spatial task either one or five black dots were presented randomly across 52 possible locations arranged on four invisible circles around fixation. After a 3000 ms delay a probe appeared in the form of a circle that matched or did not match the location of one of the previously displayed dots. Participants needed to indicate whether the location of the probe matched the location of the dot on the screen. As in the verbal task, the control condition only had a 100 ms delay and all but one circle were greyed out. Locations were not repeated over three consecutive trials. Control tasks were designed to match experimental conditions on as many elements as possible (motor response, decision making, visual stimuli, luminance) except for the working memory component. For both, verbal and spatial conditions, probes matched the target 50% of the time. The total duration of the experimental and control trials were matched. For each task, a total of 64 trials (32 experimental, 32 control) were presented in 16 blocks of four trials. Blocks alternated between experimental and control conditions in a pseudorandom fashion. Each condition was repeated four times. Total trial length was 5400 ms, which equates to a block length of 21.6 s together with an inter-block interval of 3000 ms resulting in a total scan time of 7 min and 10 s for each task. For the purpose of the current analysis we combined high and low load conditions to maximize the number of trials to be used in the MVPA analysis. The task was programmed and presented using Presentation Software Version 14.2 (Neurobehavioral Systems, Berkeley). Stimuli were displayed using Helvetica font (size: 45 pt) in black on a white background. The order of presentation of verbal and spatial tasks in the scanner was counterbalanced across participants. All young people were allowed to practice sample trials of the task on a Toshiba laptop before entering the scanner. The scripts for the tasks can be downloaded from our OSF repository: https://osf.io/a5349/.

### Results

**Table S1:** Mean (SD) for task performance measures for each group for the verbal and spatial task. Accuracy refers to the number of correct trials (out of 28 total trials per experimental and control condition), reaction times are presented in milliseconds (ms).

| | **Accuracy (Number of Correct Trials)** | | | | **Reaction Times (ms)** | | | |
|---|---|---|---|---|---|---|---|---|
| | **Experimental** | | **Control** | | **Experimental** | | **Control** | |
| | **Spatial** | **Verbal** | **Spatial** | **Verbal** | **Spatial** | **Verbal** | **Spatial** | **Verbal** |
| **TD** | 25.22 | 25.53 | 30.67 | 31.00 | 1112 | 1059 | 914 | 872 |
| **(n=19)** | (2.90) | (3.86) | (1.75) | (1.33) | (328) | (253) | (308) | (301) |
| **PDD** | 23.62 | 23.19 | 29.94 | 30.56 | 1305 | 1132 | 906 | 823 |
| **(n=16)** | (4.54) | (2.64) | (2.29) | (2.10) | (659) | (472) | (331) | (205) |
| **ADHD** | 23.95 | 23.40 | 30.20 | 29.95 | 1157 | 1043 | 892 | 853 |
| **(n=20)** | (4.41) | (3.59) | (2.02) | (2.26) | (544) | (190) | (170) | (191) |

*Head Motion*

To test for effects of head motion between the task conditions within each group, we conducted t-tests for the Euclidean norm of the motion parameters (enorm) between spatial vs. verbal stimulus material in both working memory conditions and control conditions. In the working memory conditions, head motion did not differ between the verbal and spatial working memory task components (PDD $t(15)=1.70$, $p=0.110$; ADHD $t(19)=-.02$, $p=0.981$; TD $t(18)=-.23$, $p=0.822$). In the control conditions, head motion also did not differ between the two tasks (PDD $t(15)=1.45$, $p=0.167$; ADHD $t(19)=0.44$, $p=0.665$; TD $t(18)=-1.31$, $p=0.207$). All means and standard deviations of the enorm by condition are listed in Table S2.

**Table S2:** Mean (SD) of the Euclidean norm of head motion for the verbal and spatial task for each group.

| | Experimental | | Control | |
|---|---|---|---|---|
| | **Spatial** | **Verbal** | **Spatial** | **Verbal** |
| **TD (n=19)** | .119 (.227) | .124 (.254) | .110 (.119) | .124 (.230) |
| **PDD (n=16)** | .157 (.275) | .119 (.167) | .161 (.254) | .135 (.169) |
| **ADHD (n=20)** | .208 (.361) | .205 (.358) | .210 (.376) | .200 (.342) |

*Univariate Results*

Figure S1 presents the verbal and spatial beta estimates for the experimental condition for each group. These images were fed into the MVPA analysis for classification. In accordance with predictions, the tasks significantly activate regions in the frontoparietal network and deactivate midline regions associated with the DMN in each group. Peak voxels for each cluster are shown in Table S3.

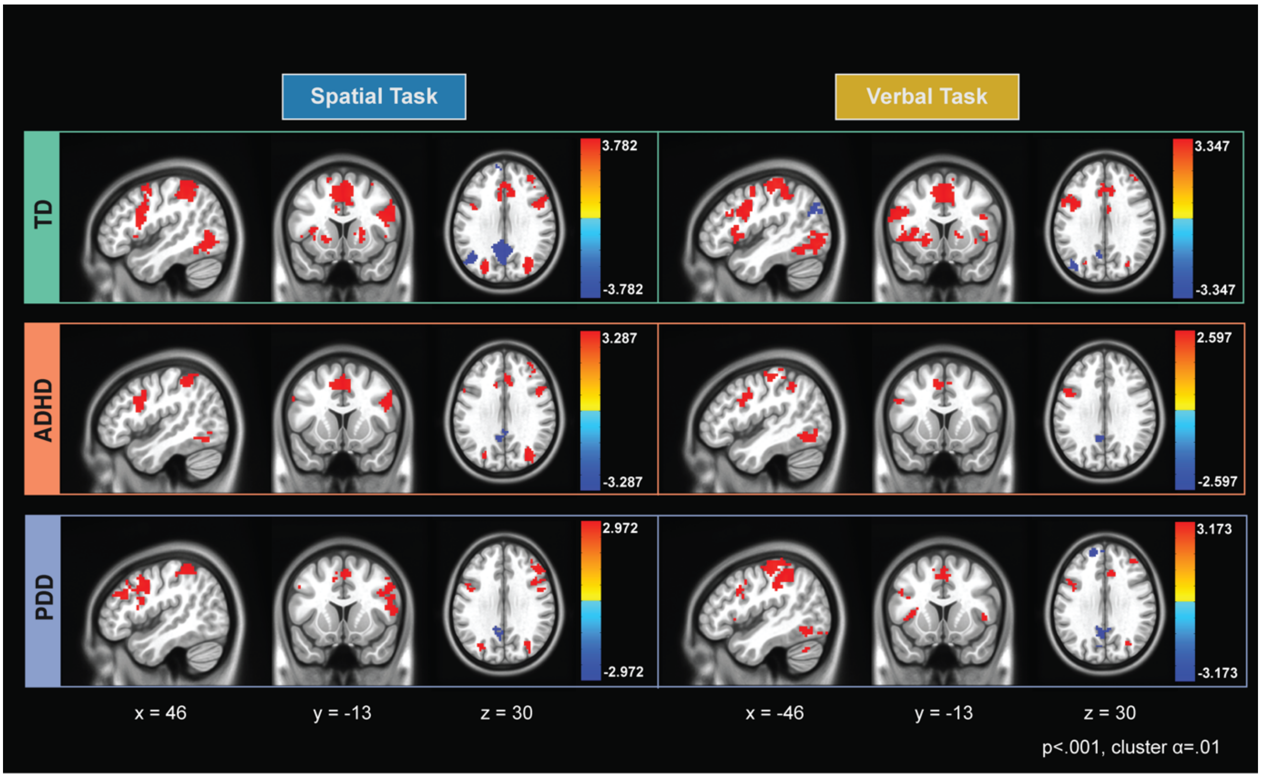

**Figure S1:** Whole-Brain univariate results for the spatial and verbal working memory experimental condition in typical development, ADHD, and PDD groups. These results revealed all groups show expected activation in key working memory related networks (e.g., left and right frontoparietal networks, salience network) and expected deactivation in default mode network for both spatial and verbal tasks. Importantly, these data show consistent activation in these networks across all groups for both spatial and verbal tasks, which markedly deviate from the neural profiles of clinical groups that were established from the MVPA decoding analyses. Signification clusters were thresholded at $p<.001$ and cluster corrected with a threshold of $\alpha = .01$. The colors correspond to t-values, with yellow-red associated with positive values and blues associated with negative values.

**Table S3:** MNI Coordinates of significant peak activations for both spatial and verbal working memory tasks. Signification clusters were thresholded at p<.001 and cluster corrected with a threshold of α = .01. Asterisk(*) denotes significant negative activation.

| Contrast | Cluster | Voxels | X | Y | Z | Stat |
|---|---|---|---|---|---|---|
| ***TD*** | | | | | | |
| *Verbal Task* | | | | | | |
| Positive | Dorsal Anterior Cingulate Cortex | 593 | 5 | -11 | 53 | 6.61 |
| | Left Anterior Insula | 416 | 44 | -20 | 8 | 5.96 |
| | Left Precentral Cortex | 385 | 38 | 29 | 62 | 6.09 |
| | Right Inferior Temporal Cortex | 376 | -44 | 68 | -2 | 5.04 |
| | Left Inferior Temporal Cortex | 300 | 47 | 77 | -2 | 5.28 |
| | Left Superior Parietal Lobule | 222 | 26 | 74 | 44 | 5.49 |
| | Left Dorsolateral Prefrontal Cortex | 218 | 44 | -11 | 32 | 4.98 |
| | Right Anterior Insula | 169 | -35 | -26 | 2 | 5.31 |
| | Right Superior Parietal Lobule | 129 | -32 | 65 | 56 | 4.77 |
| | Right Cerebellum | 85 | -11 | 71 | -23 | 5.77 |
| Negative | Left Posterior Cingulate* | 164 | 14 | 59 | 17 | -5.53 |
| | Ventromedial Frontal Cortex* | 113 | 8 | -56 | -2 | -5.33 |
| | Left Precuneus* | 64 | 41 | 77 | 35 | -4.54 |
| | Left Parahippocampal Cortex* | 41 | 32 | 41 | -8 | -4.89 |
| *Spatial Task* | | | | | | |
| Positive | Bilateral Inferior Parietal Cortex | 1986 | 44 | 44 | 41 | 6.15 |
| | Dorsal Anterior Cingulate Cortex | 109 | -2 | -23 | 41 | 6.18 |
| | Left Inferior Temporal Cortex | 440 | 47 | 74 | -5 | 5.40 |
| | Right Inferior Temporal Cortex | 182 | -50 | 59 | -11 | 4.79 |
| | Left Precentral Cortex | 175 | 32 | 5 | 56 | 5.86 |
| | Left Anterior Insula | 146 | 32 | -26 | -5 | 5.74 |
| | Right Anterior Insula | 136 | -32 | -26 | -5 | 6.46 |
| | Left Cerebellum | 128 | 8 | 74 | -20 | 5.64 |
| | Right Cerebellum | 74 | -32 | 50 | -32 | 5.07 |
| | Right Middle Frontal Gyrus | 65 | -35 | -44 | 32 | 4.47 |
| | Left Thalamus | 54 | 11 | 17 | 8 | 5.26 |
| | Left Putamen | 40 | 23 | -14 | -2 | 4.83 |
| | Left Cuneus | 33 | 14 | 71 | 5 | 3.83 |
| | Left Dorsolateral Prefrontal Cortex | 30 | 47 | -8 | 29 | 4.05 |
| | Left Caudate | 29 | 17 | 2 | 17 | 4.39 |
| | Right Caudate | 28 | -14 | -20 | 2 | 4.04 |
| Negative | Ventromedial Frontal Cortex* | 532 | 5 | -53 | -14 | -5.50 |
| | Posterior Cingulate Cortex* | 515 | 8 | 53 | 32 | -5.86 |
| | Left Angular Gyrus* | 164 | 47 | 68 | 32 | -5.30 |
| | Left Orbitofrontal Cortex* | 66 | 38 | -35 | -11 | -4.62 |
| | Left Inferior Frontal Cortex* | 53 | 56 | -32 | 2 | -4.80 |
| | Left Superior Frontal Cortex* | 41 | 14 | -47 | 47 | -4.78 |
| ***ADHD*** | | | | | | |
| *Verbal Task* | | | | | | |
| Positive | Left Precentral Cortex | 160 | 35 | 23 | 68 | 4.87 |
| | Dorsal Anterior Cingulate Cortex | 131 | 8 | -11 | 50 | 4.89 |
| | Left Inferior Temporal Cortex | 85 | 44 | 65 | -11 | 5.64 |
| | Left Dorsolateral Prefrontal Cortex | 64 | 53 | -11 | 32 | 4.67 |
| | Left Inferior Parietal Cortex | 32 | 47 | 47 | 47 | 4.27 |
| | Right Inferior Temporal Cortex | 24 | -50 | 77 | -8 | 4.25 |
| Negative | Left Posterior Cingulate Cortex* | 161 | 8 | 53 | 32 | -5.68 |
| | Right Posterior Cingulate Cortex* | 80 | -14 | 59 | 20 | -5.08 |
| | Left Parahippocampal Cortex* | 31 | 29 | 44 | -8 | -4.49 |

| | | | | | | |
|---|---|---|---|---|---|---|
| *Spatial Task* | | | | | | |
| Positive | Bilateral Inferior Parietal Cortex | 1851 | 38 | 26 | 59 | 5.73 |
| | Dorsal Anterior Cingulate Cortex | 376 | 5 | -11 | 50 | 5.40 |
| | Right Middle Frontal Gyrus | 90 | -29 | 2 | 62 | 4.41 |
| | Right Dorsolateral Prefrontal cortex | 86 | -47 | -14 | 32 | 4.91 |
| | Right Inferior Temporal Cortex | 71 | -56 | 62 | -11 | 4.44 |
| | Left Inferior Temporal Cortex | 71 | 53 | 68 | -5 | 4.72 |
| | Left Precentral Cortex | 59 | 44 | 2 | 38 | 4.71 |
| | Left Anterior Insula | 57 | 32 | -26 | 5 | 4.91 |
| | Right Dorsolateral Prefrontal Cortex | 54 | -41 | -38 | 35 | 4.69 |
| | Right Anterior Insula | 50 | -35 | -26 | -2 | 5.61 |
| | Left Cerebellum | 30 | 11 | 77 | -26 | 4.34 |
| | Right Visual Cortex | 26 | -32 | 95 | 5 | 4.10 |
| Negative | Ventromedial Frontal Cortex* | 129 | -2 | -56 | -8 | -4.77 |
| | Left Posterior Cingulate Cortex* | 55 | 5 | 50 | 14 | -4.74 |
| | Left Posterior Cingulate Cortex* | 44 | -2 | 44 | 32 | -4.51 |
| ***PDD*** | | | | | | |
| *Verbal Task* | | | | | | |
| Positive | Left Inferior Parietal Cortex | 419 | 29 | 56 | 47 | 4.99 |
| | Dorsal Anterior Cingulate Cortex | 246 | -2 | -20 | 47 | 4.82 |
| | Right Inferior Parietal Cortex | 156 | -32 | 71 | 38 | 3.80 |
| | Left Inferior Temporal Cortex | 138 | 41 | 65 | -11 | 4.83 |
| | Left Anterior Insula | 97 | 35 | -20 | 11 | 5.34 |
| | Right Anterior Insula | 91 | -35 | -23 | 8 | 4.90 |
| | Right Inferior Temporal Cortex | 86 | -50 | 68 | 2 | 4.72 |
| | Left Cerebellum | 76 | 38 | 71 | -32 | 4.78 |
| | Right Cerebellum | 71 | -29 | 68 | -26 | 5.11 |
| | Left Dorsolateral Prefrontal Cortex | 61 | 50 | -5 | 38 | 4.77 |
| | Right Dorsolateral Prefrontal Cortex | 45 | -41 | -47 | 20 | 4.04 |
| | Right Occipital Cortex | 42 | -41 | 92 | -2 | 4.65 |
| | Mid Cerebellum | 31 | -11 | 71 | -26 | 4.90 |
| | Left Thalamus | 28 | 14 | 14 | 5 | 4.63 |
| Negative | Posterior Cingulate Cortex* | 321 | -8 | 53 | 20 | -5.10 |
| | Ventromedial Prefrontal Cortex* | 174 | 2 | -41 | -5 | -4.79 |
| | Left Superior Frontal Cortex* | 74 | 20 | -50 | 41 | -4.62 |
| | Right Orbitofrontal Cortex* | 28 | -38 | -38 | -11 | -4.59 |
| | Right Superior Frontal Cortex* | 28 | -23 | -35 | 44 | -4.61 |
| | Left Parahippocampal Cortex* | 26 | 32 | 35 | -14 | -4.32 |
| | Right Parahippocampal Cortex* | 24 | -26 | 45 | -14 | -4.08 |
| *Spatial Task* | | | | | | |
| Positive | Left Inferior Parietal Cortex | 521 | 50 | 44 | 53 | 5.80 |
| | Right Inferior Parietal Cortex | 436 | -38 | 56 | 56 | 5.69 |
| | Right Dorsolateral Prefrontal Cortex | 273 | -50 | -11 | 41 | 4.76 |
| | Dorsal Anterior Cingulate Cortex | 148 | -5 | -11 | 53 | 4.98 |
| | Left Precentral Cortex | 90 | 29 | 5 | 62 | 5.19 |
| | Left Precentral Cortex | 50 | 50 | -5 | 38 | 4.90 |
| | Right Superior Frontal Cortex | 45 | -26 | -2 | 59 | 4.16 |
| Negative | Left Posterior Cingulate Cortex* | 65 | 11 | 50 | 32 | -3.98 |

*NN1 for activation p < .001 for all images, cluster corrected at p<.01*
